# Energy expenditure and body composition changes after an isocaloric ketogenic diet in overweight and obese men: a secondary analysis of energy expenditure and physical activity

**DOI:** 10.1101/383752

**Authors:** Mark I. Friedman, Scott Appel

## Abstract

**Background:** A previously published pilot study assessed energy expenditure (EE) of participants with overweight and obesity after they were switched from a baseline high-carbohydrate diet (BD) to an isocaloric low-carbohydrate ketogenic diet (KD). EE measured using metabolic chambers increased transiently by what was considered a relatively small extent after the switch to the KD, whereas EE measured using doubly labeled water (EE_DLW_) increased to a greater degree after the response in the chambers had waned. Using a publicly available dataset, we examined the effect of housing conditions on the magnitude of the increase in EEDLW after the switch to the KD and the role of physical activity in that response.

**Methods:** The 14-day EE_DLW_ measurement period included 4 days when subjects were confined to chambers instead of living in wards. To determine the effect on EE_DLW_ only for the days subjects were living in the wards, we calculated non-chamber EE (EE_nonchamber_). To assess the role of physical activity in the response to the KD, we analyzed chamber and non-chamber accelerometer data for the BD and KD EE_DLW_ measurement periods.

**Results:** In comparison with the increase in average 14-day EE_DLW_ of 151 kcal/d ± 63 (*P* = 0.03) after the switch to the KD, EE_nonchamber_ increased by 203 ± 89 kcal/d (*P* = 0.04) or 283 ± 116 kcal/d(*P* = 0.03) depending on the analytical approach. Hip accelerometer counts decreased significantly (*P* = 0.01) after the switch to the KD, whereas wrist and ankle accelerometer counts did not change.

**Conclusions:** **S**witching from the BD to the KD substantially increased EE_DLW_, but apparently only on days subjects were living in the ward outside the metabolic chamber. Increased physical activity as measured by accelerometry did not appear to account for this effect.

## Introduction

Debate over the role of diet in the etiology of obesity often centers on the relative importance of dietary macronutrient composition versus total energy consumption. According to one view of obesity [1–3], the primary cause of fat accumulation involves a shift in the partitioning of metabolic fuels away from pathways of oxidation toward those of fat synthesis and storage. In this case, diet composition can be an important contributing factor; diets rich in carbohydrate, in particular those containing substantial amounts of refined grains and sugars, promote obesity because stimulation of insulin secretion by these nutrients drives metabolic fuels toward the synthesis and storage of fat [4]. This is known as the “carbohydrate-insulin” hypothesis. The more conventional model sees obesity as caused by an energy balance disorder in which energy intake exceeds energy expenditure [5,6]. According to this “energy balance” hypothesis, this excessive total energy intake, regardless of the macronutrient source of the energy, is the primary cause of obesity.

The carbohydrate-insulin and energy balance hypotheses make distinctly different predictions about the effects of reducing dietary carbohydrate content on energy expenditure (EE) under conditions in which calories and protein remain constant. The carbohydrate-insulin hypothesis predicts that lowering the proportion of carbohydrate to fat, even while maintaining energy and protein intake, would minimize circulating insulin concentration and thereby promote lipolysis and oxidation of stored and ingested fat, and, as a result, increase EE. On the other hand, the energy balance hypothesis, which assumes “a calorie is a calorie,” predicts that exchanging fat calories for carbohydrate calories would have no appreciable effect on energy expenditure [7].

In preparation for an anticipated full-scale trial to test these competing predictions, Hall et al. conducted a pilot study [8,9] in which they measured EE in participants with overweight and obesity who were housed in metabolic wards before and after they were switched from a high-carbohydrate baseline diet to an isocaloric ketogenic diet containing equivalent protein and little carbohydrate. EE was measured two ways: using metabolic chambers for two consecutive days each week throughout the study, and using doubly labeled water during the last 2 weeks of each 4-week diet period. EE measured in metabolic chambers increased significantly after the switch to the ketogenic diet, but this change was transient, lasting only two weeks, and was considered to be relatively small by Hall et al., which led them to conclude that the results did not support the carbohydrate-insulin model. However, in contrast to the results using metabolic chambers, EE, measured using doubly labeled water (EE_DLW_) after the response in the chambers had waned, increased more substantially after the switch to the ketogenic diet (151 kcal/d vs. 57 kcal/day). This response was attributed [8] to greater energy expenditure from increased physical activity when subjects ate the ketogenic diet and were outside the chambers living in the ward.

The carbohydrate-insulin and energy balance hypotheses have distinctly different implications for understanding the etiology of obesity and devising effective strategies for preventing and treating it. Consequently, it would be useful to reconcile the discrepant findings from measurements of EE using metabolic chambers and doubly labeled water. Hall et al. made the data from their study publicly available on the Open Science Framework (OSF) website [10]. In this paper, we report results of additional analyses of this dataset to differentiate the effect of housing subjects in a metabolic chamber versus in a metabolic ward on the magnitude of the increase in EE_DLW_ after the switch to the ketogenic diet and to assess the role of physical activity in this effect.

## Methods

### Overview of the Hall et al. study

Details of the design and methods of the study can be found in the Hall et al. paper [8], including the online supplementary data [11], and in the published IRB-approved protocol [9]. Briefly, focusing on methods relevant to the analyses described here, 17 males with overweight or obesity were admitted as inpatients to metabolic wards and fed a baseline diet (BD; 15:50:35 percent of calories from protein:carbohydrate:fat) for 4 weeks followed by an isocaloric ketogenic diet (KD; 15:5:80 percent of calories from protein:carbohydrate:fat) for another 4 weeks. Subjects were housed in a metabolic chamber for two consecutive days each week throughout the study to measure daily EE, sleeping EE, and respiratory quotient (considered primary endpoints of the study). During the last 2 weeks of each diet period, average daily EE was measured using doubly labeled water (EE_DLW_; considered an exploratory endpoint of the study). Physical activity level was monitored throughout the study using accelerometers; each subject wore an accelerometer (GT3XE+; Actigraph Corporation) on a hip, wrist and ankle, and accelerometer counts and the length of time wearing the devices were logged for each device location. The study was registered at www.clinicaltrials.gov as NCT01967563.

### Reproducing calorimetry results

To confirm the replicability of the data used in the secondary analyses of EE_DLW_ described below, we first reanalyzed the calorimetry results reported in Table 2 of the Hall et al. paper [8] using the dataset and code published on the OSF website [10] and SAS v9.4 (SAS Institute, Inc.).

Details regarding the sources and handling of data from the Hall et al. dataset for the secondary analyses described below are provided in the Supporting Information (S1 File) along with the SAS code used for these secondary analyses (S2 File). All endpoint values reported herein were calculated using individual data from the Hall et al. dataset.

### Non-chamber EE_DLW_

A primary purpose of the Hall et al. pilot study was to determine the magnitude and variability of changes in EE after subjects were switched from the BD to KD in preparation for an anticipated larger study. The 14-day period for measuring EE_DLW_ included 4 days when subjects were confined to a metabolic chamber and 10 days when subjects lived in the ward. Hall et al. reported EE_DLW_ as a daily average across the 14-day measurement period and did not differentiate EE during the non-chamber days, when subjects were housed in the ward, from the chamber days, when EE is relatively lower [12,13] and the effect of diet was much reduced [8].

To determine average daily EE_DLW_ for only those days in which subjects were housed in the ward, we used a term in Hall et al.’s Equation 6 for calculating non-chamber EE (EE_nonchamber_) [8]. In essence, the resulting equation (Equation 1 below) separates average daily EE for days subjects were housed in the wards from days they were confined to metabolic chambers by subtracting total EE measured during the 4 chamber days (EE_chamber_) within the EE_DLW_ measurement period from total 14-day EE_DLW_ and averaging the resulting value over the 10 non-chamber days.

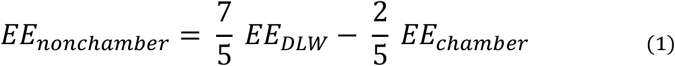

Differences in EE_DLW_, EE_nonchamber_ and EE_chamber_ between the two diet conditions were evaluated using a repeated linear mixed model. Data from Subject 04-012 was not included in these analyses (see below) in keeping with Hall et al. [8]. A *P* value of < 0.05 (two-sided tests) was considered statistically significant for this and all other analyses below.

As a check on Equation 1, we also calculated EE_nonchamber_ by subtracting total CO_2_ production measured during the four chamber days within the EE_DLW_ measurement periods from the total 14-day CO_2_ production measured using doubly labeled water, and dividing the result by the 10 non-chamber days. The resulting *r*CO_2_ values for the BD and KD conditions were converted to kcal/d using equations described in Hall et al. (8). Details of these calculations are provided in the Supporting Information (S1 file).

### EE_DLW_ outlier

Hall et al. excluded one subject’s (#04-012) data from the analysis of energy expenditure measured using doubly-labeled water. This subject showed the largest increase in EE_DLW_ after the switch from the BD to the KD (1136 kcal/d), which was identified statistically as an outlier value using Cook’s distance. Because this subject’s relatively extreme change in EE_DLW_ was not apparently due to a documented error in, for example, data collection, recording, computation or coding, best practices [14] indicate that the EE_DLW_ analysis should be reported with and without the outlier data. To that end, we compared EE_DLW_ and EE_nonchamber_ during the two diet periods, as above, except that data from Subject 04-012 were included in the analysis.

Hall et al. justified exclusion of Subject 04-012’s data on the basis that he gained 0.2 kg during the KD period despite the marked increase in EE_DLW_ after the switch to the KD and an EE_DLW_ during the KD period that substantially exceeded his energy intake. To determine whether the extreme change in Subject 04-012’s EE_DLW_ was reflected in other parameters related to his energy balance in the KD period, we examined his change in body weight, absolute EE_DLW_ and energy intake, and the difference between EE_DLW_ and intake relative to the entire group of subjects during the KD EE_DLW_ measurement period.

The reported 0.2 kg weight gain occurred over two body composition assessments performed during the latter part of the KD period. As a check on the weight change based on body weights collected during the body composition assessments, we evaluated the change in body weight during the EE_DLW_ measurement period using daily body weight data from the dataset.

### Accelerometer data

Hall et al. evaluated whether greater physical activity accounted for the increase in EE_DLW_ during the KD period by calculating energy expenditure from physical activity in and out of the metabolic chambers (i.e, PAE_chamber_ and PAE_nonchamber_ as per their Table 2). Physical activity energy expenditure outside the chambers was higher during the KD period compared to the BD phase, but the effect was not statistically significant. Physical activity level was measured directly throughout the study using accelerometers; however, only hip count data were reported and only as a percentage difference between chamber and non-chamber days during the entire BD period. Here, we used the accelerometer data in the OSF dataset to determine more directly whether differences in physical activity can account for the increase in EE_DLW_ after the switch to the KD.

To confirm reproducibility and help validate our use of the accelerometer data, we first reanalyzed the fractional difference between hip accelerometer counts from chamber and non-chamber days during the baseline period of the study using the published dataset and code. We next analyzed daily hip, wrist and ankle accelerometer counts during the BD and KD EE_DLW_ measurement periods with respect to whether subjects were confined to chambers or were housed in the ward (i.e., chamber and non-chamber days) using generalized linear mixed models. Average accelerometer wear times varied little with respect to device location, diet and housing status. In keeping with Hall et al.’s calculation and analysis of physical activity energy expenditure, accelerometer data were analyzed excluding data from Subject 04-012. In a separate analysis, this subject’s data was included. Accelerometer counts for each location with respect to chamber status and diet were compared in the generalized linear mixed model by t-test to determine statistical significance.

### Sleeping energy expenditure

Hall et al. reported that sleeping energy expenditure (SEE; kcal/d) measured in the metabolic chambers increased in the first week after subjects were switched from the BD to the KD, and then declined during the subsequent three weeks. To determine whether this increase in SEE persisted during the end of the KD period and may have contributed to the increase in EE_DLW_ observed at that time, we compared SEE during the BD and KD EE_DLW_ measurement periods. Comparison of SEE for all subjects during the EE_DLW_ measurement periods in the BD and KD phases of the study were made using a paired t-test.

## RESULTS

### Reproducing calorimetry results

Reanalysis of the calorimetry data in Table 2 in the Hall et al. paper [8] using the OSF dataset and code fully reproduced the reported results, including mean, standard error and probability values associated with statistical comparisons of diet periods.

### Non-chamber EE_DLW_

The increase in EE_DLW_ after the switch to the KD was greater when calculated only for days when subjects were housed in the wards outside of the chambers than it was when calculated over the entire EE_DLW_ measurement period that included both days in and out of the chambers. Reanalysis of EE_DLW_ data from the Hall et al. dataset reproduced the average 151 kcal/d increase in EE_DLW_ after subjects were switched to the KD (Table 1). Using Equation 1 above, energy expenditures for days when subjects were out of the chambers (EE_nonchamber_) increased on average by 203 kcal/d after subjects were switch from the BD to the KD. Energy expenditure measured in the chambers (EE_chamber_) during the EE_DLW_ measurement periods did not differ as a function of diet.

**Table1.** Energy expenditure during the BD and KD doubly labeled water measurement periods^a^

|  | **BD** | **KD** | **Difference** | ***P*<sup>b</sup>** |
| --- | --- | --- | --- | --- |
| **$EE_{DLW}$** | $2995 \pm 45$ | $3146 \pm 45$ | $151 \pm 63$ | 0.03 |
| **$EE_{nonchamber}$** | $3142 \pm 45$ | $3344 \pm 45$ | $203 \pm 89$ | 0.04 |
| **$EE_{chamber}$** | $2628 \pm 22$ | $2649 \pm 22$ | $21 \pm 32$ | 0.52 |
| **$EE_{DLW}$ (*included*)** | $2964 \pm 59$ | $3173 \pm 59$ | $209 \pm 83$ | 0.02 |
| **$EE_{nonchamber}$ (*included*)** | $3100 \pm 82$ | $3382 \pm 82$ | $283 \pm 116$ | 0.03 |

<sup>a</sup>Data are least squares mean ± SEs from a linear mixed model. Values are based on $n = 16$ except for those designated as “*included*” ( $n = 17$ ), which include an outlier removed from the other analyses as described in the text. Difference values are the change from the BD to KD period. $EE_{chamber}$ , daily energy expenditure measured for 4 days in metabolic chambers during the 14-day doubly labeled water measurements; $EE_{DLW}$ , average energy expenditure over 14 days measured using doubly labeled water; $EE_{nonchamber}$ , average energy expenditure over 10 days outside metabolic chambers.
<sup>b</sup>Values refer to the difference between diet periods by t-test using modeled standard errors.

Calculation of EE_nonchamber_ based on the difference between CO_2_ production measured in the chambers and by doubly labeled water produced results very similar to those using Equation 1. With Subject #04-012 removed from the analysis, EE_nonchamber_ during the BD and KD periods were, respectively, 3140 ± 146 kcal/d and 3355 ± 189 kcal/d with a significant difference of 215 ± 87 kcal/d (*P* = 0.03 by paired t-test).

### EE_DLW_ outlier

The effect of switching from the BD to the KD on EE measured using doubly labeled water was greater when data from the outlier was included in the analysis. As shown in Table 1, the increase in EE_DLW_ after the switch to the KD was greater when Subject 04-012’s data were included in the analysis than when they were not (209 ± 83 kcal/d vs. 151 ± 63 kcal/d). When this subject’s data were included in the calculation of EE_nonchamber_, the effect of switching to the KD was greater (283 + 116 kcal/d) than when his data were excluded (203 ± 89 kcal/d; see Table 1). With all subjects included in the analysis, EE_nonchamber_ during the BD and KD periods, as calculated using CO_2_, were, respectively, 3100 ± 142 kcal/d and 3394 ± 182 kcal/d with a significant difference of 295 ± 113 kcal/d (*P* = 0.02 by paired t-test).

Subject 04-012’s weight gain, absolute EE_DLW_ and energy intake, and the difference between his EE_DLW_ and intake during the KD EE_DLW_ measurement period as reported by Hall et al., were each within the variability of the group as a whole. The 0.2 kg weight gain of Subject 04-012 was within one standard deviation of the average weight change of all subjects (−0.6 kg + 0.8, mean ± SD). His EE_DLW_ (3612 kcal/d) and energy intake (2794 kcal/d), and difference between them (818 kcal/day) were well within one standard deviation from the means of the group (3173 ± 583 kcal/day, 2736 ± 428 kcal/day, and 437 ± 481 kcal/day, respectively; mean ± SD).

Inspection of the dataset revealed that other subjects showed changes in body weight that appeared anomalous relative to the difference between their EE_DLW_ and energy intakes. Two of these subjects gained weight during the KD period (0.6 and 1.3 kg) despite a difference in expenditure and intake of, respectively, 1751 and 465 kcal/d. Two participants lost weight (1.9 and 0.2 kg) despite an excess of energy intake relative to EE_DLW_ of (291 and 250 kcal/d, respectively.

The weight gain of Subject 04-012 across the interval between two body composition assessments in the KD period reported by Hall et al. underlies their rationale for exclusion of his data from analysis of the effect of diet on EE_DLW_. We confirmed that Subject 04-012 gained 0.2 kg between the two body composition assessments during the KD period; however, inspection of the dataset also revealed that the interval between the two body composition assessments and the EE_DLW_ measurement period were not concurrent. Consequently, we referred to daily body weight data from the dataset, which showed that Subject 04-012 lost 0.5 kg over the EE_DLW_ measurement period.

### Accelerometer data

Reanalysis of hip accelerometer counts during the full BD period using the Hall et al. code reproduced their finding that counts were 21 ± 4% greater on non-chamber days than they were on days when subjects were confined to metabolic chambers.

Although during the DLW measurement periods hip, wrist, and ankle accelerometer counts were significantly greater when subjects were housed in the ward than when they were confined to metabolic chambers (*P*’s < 0.001), counts either decreased (hip; *P* = 0.006) or did not change significantly (wrist and ankle) after the switch from the BD to the KD (Table 2). Inclusion of data from Subject 04-012 did not materially affect accelerometer counts or the outcomes of the statistical analyses.

**Table 2.**
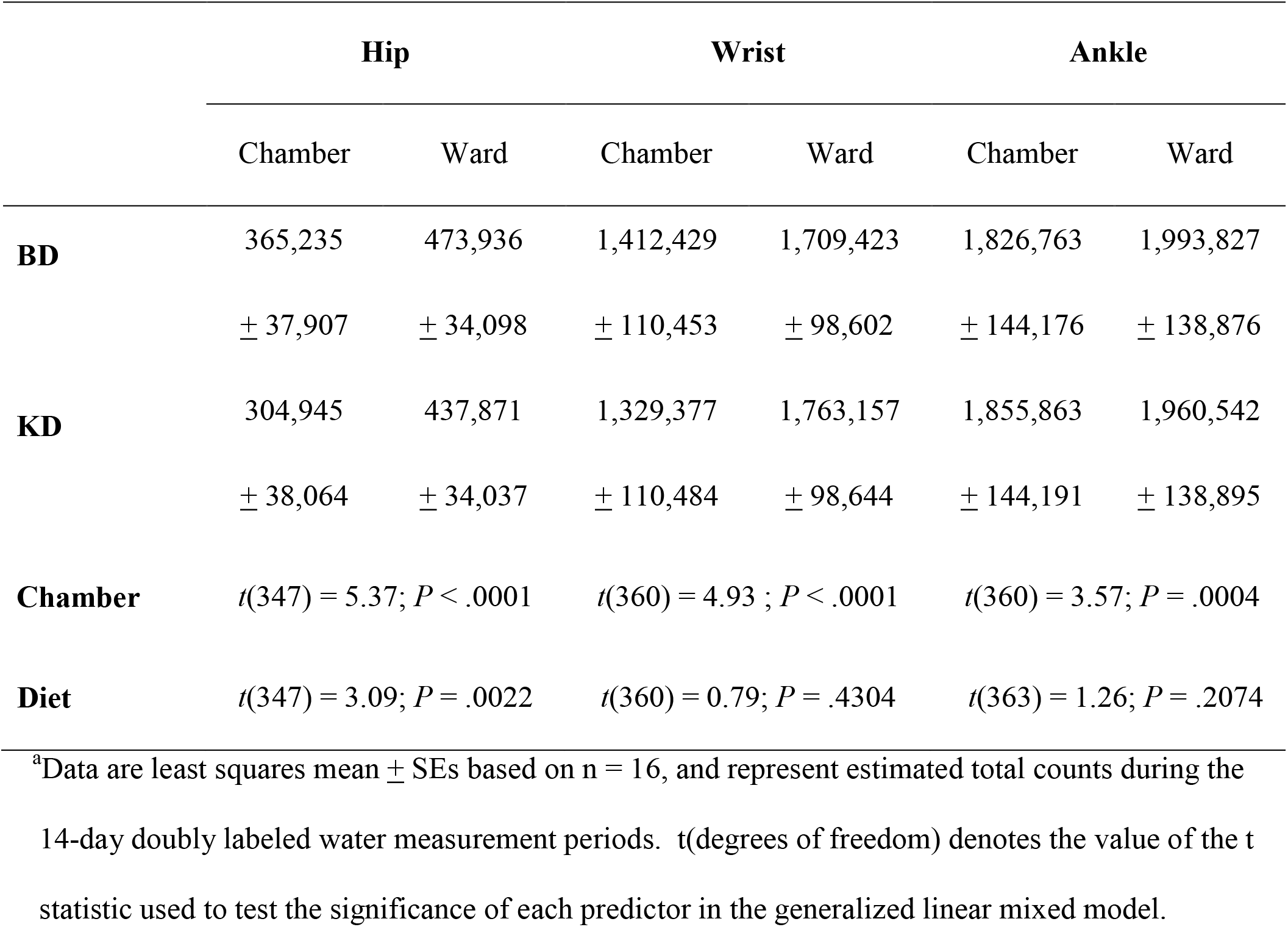
Accelerometer counts during the BD and KD doubly labeled water measurement periods as a function of housing condition^a^

Interactions between chamber status and diet were 269 not significant (*P* = 0.40, 0.20, and 0.35 for hip, 270 wrist, and ankle, respectively).

### Sleeping energy expenditure

SEE measured in the chambers during EE_DLW_ measurement periods was significantly greater after the switch from the BD to the KD (BD, 1576 ± 68 kcal/d and KD 1620 ± 56 kcal/d; difference = 44 ± 18 kcal/d, *P* < 0.02). SEE values and the results of the analysis were nearly identical if data from Subject 04-012 were excluded.

## Discussion

### Non-chamber energy expenditure – effect size

Hall et al. measured average daily energy expenditure using doubly labeled water over a 14-day period that included 4 days during which subjects were confined to a metabolic chamber and 10 days when they were housed in the ward. Because people expend less energy in a metabolic chamber than under more free-living conditions [12,13] and chamber measurement of EE showed little difference between the diet periods in the Hall et al. study [8], we quantified expenditures for non-chamber days (EE_nonchamber_) separately from in-chamber days (EE_chamber_). The results showed that the switch from the BD to the KD diet was accompanied by an increase in EE_DLW_ that was 34% greater than that originally reported [8]. EE_chamber_ during the doubly labeled water measurement periods did not differ significantly as a function of diet, further indicating that the increase in EE_DLW_ after the switch to the KD was limited to days when subjects were housed in the ward.

In keeping with best practices for handling outliers [14], we calculated EE_DLW_ and EE_nonchamber_ with and without data from Subject 04-012, considered an outlier by Hall et al. [8]. Relative to the increase in EE_DLW_ after the switch to the KD as originally reported, including these data increased the effect on EE_DLW_ and EE_nonchamber_ by, respectively, 38% and 87%. Although, as discussed below, exclusion of this subject’s data seems unwarranted based on changes in his body weight relative to expenditures, his outlier status regarding the increase in EE_DLW_ from the BD to KD period is clear on a statistical basis. However, because there is no documented error in, for example, data collection, recording or computation, the cause of this apparently exaggerated response is unknown. Taken together with the relatively small number of subjects in this pilot study, and the lack of previous research to help determine the range of response to KD diets under these controlled conditions, additional research will be required to determine whether this subject’s data are invalid or reflect a relatively extreme response seen in a small proportion of the population.

According to Hall et al., the carbohydrate-insulin model predicts that consuming a KD would increase energy expenditures by 300-600 kcal/d. The higher value is suspect, having been based on a theoretical estimate [15] of a 400-600 kcal/d expenditure to support gluconeogenesis under conditions very different than those in the Hall et al. study, specifically prior to adaptation to a low-carbohydrate diet with only endogenous, not dietary, protein as the substrate. In contrast, Bistrian [16] recently estimated the energy cost of gluconeogenesis associated with consumption of a ketogenic diet at 110 kcal/day (not allowing for tissue glucose demands) under conditions very similar to those employed in Hall et al. (i.e., eucaloric after an adaptation period). The lower value cited by Hall et al.’s (300 kcal/d) was based on the results of a randomized cross-over study by Ebbeling et al. [17] in which EE_DLW_ in free-living weight-reduced subjects was, depending on how expenditures were calculated, ~250-325 kcal/d greater when subjects ate a low-carbohydrate/high-fat diet compared with a high-carbohydrate/low-fat diet. More recently, Ebbeling et al. [18] reported results from a larger and longer duration randomized group trial in which EE_DLW_ in free-living weight-reduced subjects eating a low-carbohydrate diet was, depending on whether data were analyzed by an intention-to-treat or per protocol analysis, ~200-280 kcal/d greater than that of subjects consuming a high-carbohydrate diet. Whereas these effects of diet on EE_DLW_ in these two studies were observed under very different conditions than those in the Hall et al. study, including during maintenance of a 10-15% weight loss, the magnitude of the effects was similar to the increase of ~200-280 kcal/d in EE_nonchamber_ after the diet switch in the Hall et al. study.

The order in which subjects in the Hall et al. study were fed the BD and the KD was not counterbalanced or otherwise controlled for, a trial design limitation noted by the authors that precluded causal inference about the effect of the KD. In contrast, the Ebbeling et al. studies described above randomized the order in which subjects ate the experimental diets [17] or randomized the diets to which they were assigned [18]. The similarity in the responses to a low carbohydrate diet in the Ebbeling et al. studies and, with respect to EE_nonchamber_, to the KD in the Hall et al. study, lends credence to the conclusion that consumption of the KD caused the increase in EE_DLW_ in the Hall et al. study.

Estimates of the increase in EE_nonchamber_ from the current analysis likely represent a minimal range for the effect size. The continuing weight loss throughout the study due to unintentional underfeeding of the subjects, as described by Hall et al., would be expected to have suppressed EE [13], thereby mitigating any increase in energy expenditure after switching to the KD. Lower circulating concentrations of leptin and triiodothyronine during the KD versus the BD period reported by Hall et al. are consistent with such a reduction in EE. Accounting for the excretion of fat in feces might also magnify energy losses during the KD period [].

### Methodological considerations

Hall et al. [20] recently argued that the increase in EE_DLW_ after the switch to the KD in their earlier pilot study [8] was partially due to methodological issues associated with the doubly labeled water technique. Based on hypothetical relationships between diet composition, energy balance and measured RQ, they suggested that their earlier calculation of EE_DLW_ overestimated the effect of switching to the KD. Adjusting for these factors, they found that the increase in EE_DLW_ after the diet switch was diminished to statistically nonsignificant levels, an effect that was due primarily to an increase in estimated EE_DLW_ in the BD period as opposed to a change in the KD period. Hall et al. [20] identified two participants as outliers (Subjects A and B; Subject B is Subject #04-012 referred to above). Excluding these subjects’ data from the new analysis largely eliminated the effect of switching the diet on EE_DLW_. Hall et al. [8, 20] did not pre-specify criteria or methods for identifying and handling outliers. Subjects were identified as outliers *post hoc*, particularly on the basis of observations indicating that the difference between their EE_DLW_ and energy intake was not commensurate with changes in body weight during the KD EE_DLW_ measurement period.

The two outliers identified by Hall et al. [20] showed large discrepancies between EE_DLW_ and energy intake. However, as reported above in relation to analysis of Subject 04-012’s (Subject B’s) designation as an outlier and as described in the Supporting Information (S3 File, Table 1S), other subjects showed differences between expenditure and intake that, while not as large, were substantial and were also incommensurate with changes in weight. Which of the outlier data is chosen for exclusion in data analysis markedly affects estimates of energy expenditure (Supporting Information File S3, Figure 2S). In the case of Subjects A and B together, it reduced effect size to nonsignificant levels. In other cases, it reduced the effect size less while retaining statistical significance, and, importantly, in some cases the choice of outliers increased the effect size. These finding suggest that the selection of outliers by Hall et al. [20] was too restrictive and thereby may have overly circumscribed the interpretation of the results of their reanalysis.

The changes in weight that were part of the assessment of outliers in the original and recent Hall et al. studies [8, 20] were derived from body weight measurements taken over the interval between two body composition assessments in the latter part of each diet period. Hall et al. [20] claimed that assessments were coincident with the EE_DLW_ measurement periods. As discussed above, this was not the case for Subject 04-012 (Subject B), nor, for Subject A (Supporting Information File S3; Table 2S). Hall et al. reported that Subjects A and B gained weight during the EE_DLW_ measurement period based on the body composition assessment measures (0.6 and 0.2 kg, respectively). In contrast, both subjects lost 0.5 kg of weight when body weight change was measured over a period that was in fact concurrent with the EE_DLW_ measurement period using daily body weight data from the Hall et al. [8] dataset. Although the magnitude of body weight changes differed in all but one subject depending on whether it was based on body composition assessment data or was synchronized with the EE_DLW_ measurement period using daily body weight data, only Subjects A and B showed such a reversal. Differences in measurement precision does not appear to account for the difference in body weight change because weight changes across the interval for the body composition assessments measured using daily body weight data were similar to those based on weights measured during the assessments (File S3, Table 2S). Because a loss of body weight would be commensurate with an excess of energy expenditure relative to energy intake, the outlier status of Subjects A and B seems difficult to justify.

Calculation of energy balance based on changes in fat mass and fat free mass are central to Hall et al.’s reanalysis. Hall et al. [20] acknowledged uncertainly regarding these calculated values “because DXA has a limited ability to precisely and accurately detect small changes in body energy stores.” The body composition assessments were presumed to be coincident with those of the EE_DLW_ measurement period, thus providing an accurate assessment of changes in bodily energy stores associated with those in energy expenditure. However, as discussed above, the timing of body composition assessments and EE_DLW_ measurement periods differed and a difference of just a few days markedly affected estimates of changes in body weight. Such asynchrony might add additional uncertainty to estimates of body composition and, in turn, energy balance. To what extent the results of Hall et al.’s study [20] reflect the hypothetical relationships between diet composition, energy balance and RQ upon which their reanalysis is based or the imprecision of their measurements remains to be determined.

In their reconsideration of the original paper, Hall et al. [20] also suggested that higher rates of *de novo* lipogenesis during consumption of the high-carbohydrate BD as compared with the low-carbohydrate KD magnified the increase in EE_DLW_ after switching to the KD because more deuterium would be sequestered in fat during the BD period and thereby lower estimated CO_2_ production. Such an effect of deuterium trapping has been documented only in rapidly fattening piglets with no significant consequence for estimates of EE_DLW_ predicted for weight stable animals [21]. Given this, it seems unlikely that differences in *de novo* lipogenesis during the diet periods would account for much, if any, of the increase in EE_DLW_ in the original Hall et al. [8] study since subjects lost weight throughout the trial. Hall et al.’s quantitative estimates of the effects of *de novo* lipogenesis on EE_DLW_ also appear overestimated by at least two-fold. Their estimates were based on previously published total triglyceride turnover values [22], but did not take into account that *de novo* lipogenesis involves only nonessential fatty acids, which comprise only half of stored fatty acids [22]. Also, from earlier work [23] they estimated that 10-20% of the daily production of very low density lipoprotein triglycerides are derived from hepatic *de novo* lipogenesis; however, this estimate was based on meal-to-meal changes in *de novo* lipogenesis after two meals of a rapidly absorbed liquid diet [23] and did not take into account that the contribution of hepatic *de novo* lipogenesis to triglyceride production falls to < 5% during an overnight fast [24]. Clearly, a more definitive estimate of the magnitude of the effect of a ketogenic diet on EE_DLW_ will require studies using weight-stable subjects along with direct analysis of relevant biochemical and physiological processes.

A discrepancy between chamber and doubly labeled water measures of EE in response to a nutritional manipulation was reported previously by Rosenbaum et al. [13] who found substantial changes in total daily EE in response to over- and under-feeding when measured using doubly labeled water or by the non-isotopic method of caloric titration in subjects housed in a ward, but not when measured in the same subjects using chamber respirometry. These investigators hypothesized that the different outcomes between methodologies to limitations on physical activity imposed by the metabolic chamber. The present analyses of the Hall et al. data similarly suggest that the magnitude of the effect of a KD on EE depends on conditions in which physical activity is not restricted by the confines of a metabolic chamber. Although more direct methodological comparisons are needed, the preferred approach for studies of the effect of nutritional status or dietary composition on energy balance would appear to entail at least the opportunity for physical activity afforded by a metabolic ward along with appropriate methods for measurement of EE that do not require restricted confinement.

### Increased non-chamber energy expenditure – possible mechanisms

Accelerometer counts, a direct measure of physical activity, did not increase and, in the case of hip counts, decreased during the EE_DLW_ measurement period after the switch from the BD to the KD. These findings do not support the suggestion in Hall et al. [8] that the lack of a significant increase in non-chamber energy expenditure from physical activity during the KD period was due to limitations on activity imposed by the metabolic ward environment. The dissociation between changes in physical activity and EEDLW, is consistent with other findings [17,18] that free-living subjects eating a low-carbohydrate diet show elevated EE_DLW_ but little or no change in physical activity measured using accelerometers. Accelerometry in these studies and the Hall et al. study may not have captured all components of physical activity contributing to nonexercise activity thermogenesis (NEAT) [25], although the hip and wrist accelerometers counts likely reflected walking, which is a major component of NEAT [25].

Sleeping energy expenditure, derived from sleeping metabolic rate, was greater during the EE_DLW_ measurement period after the switch to the KD. Because sleeping metabolic rate approximates as much as 80-100% of basal metabolic rate [26], the difference in sleeping energy expenditure suggests that the KD may have increased basal energy expenditure by ~50 kcal/d, accounting for ~18-25% of the increase in daily energy expenditures depending on the range of estimates for the effect of the KD. Such an estimate must be tempered, however, given that physical activity during sleep will increase metabolic rate and that the difference in sleeping metabolic rate as a function of diet measured in chambers may differ from that when subjects were sleeping in the ward.

After allowing for an elevation in basal metabolic rate and some contribution from physical activity not monitored by accelerometers, much of the increase in EE_DLW_ during the KD period appears left to be explained. Given that the effect of switching to the KD is limited to days subjects were housed in the ward, the increase in EE_nonchamber_ may have been dependent on the increase in physical activity outside the chambers but not directly caused by it. Much of the increase in thermogenesis in overfed subjects housed on a metabolic ward is due to an increase in non-resting energy expenditure and about one-third of this effect has been attributed to lower skeletal muscle work efficiency [27]. This suggests that nutritional factors can affect energy expenditures and may do so in part by increasing the energetic cost of physical activity. Consumption of a very low carbohydrate, ketogenic diet could have similar effects; indeed, as early as 1920, Krogh and Lindhard [28] described the “waste” of energy from fat in exercising humans maintained on a largely fat, as compared with a primarily carbohydrate, diet. On the other hand, in the Hall et al. study [8], EE measured during the prescribed cycling exercise in the chambers was similar in the two diet periods. This observation argues against a change in muscle work efficiency in the KD period; however, it is not known whether the efficiency of muscle work associated with the increase in other forms physical activity in a ward setting may have been affected. Greater physical activity outside the chambers may also have increased EE during the KD period by creating a demand for glucose, some of which under the condition of severe dietary carbohydrate restriction would be met through the energetically expensive process of hepatic gluconeogenesis [see also 16]. The elevated plasma concentrations of glucagon and the increase in protein catabolism (as evidenced by increased urinary nitrogen, urea and ammonia excretion) during the KD period in the Hall et al. study are consistent with such a higher rate of gluconeogenesis, which has been suggested as a contributing cause of the increased thermogenesis associated with consumption of ketogenic diets [16].

## Conclusions

On the basis of the transient and what was considered a relatively small increase in EE measured in the metabolic chambers after the switch to the KD, Hall et al. [8] concluded that the results of their pilot study did not support the prediction of the carbohydrate-insulin hypothesis of obesity that such an isocaloric change in diet would increase EE. This finding has been cited as evidence refuting the carbohydrate-insulin hypothesis, offering further support for the energy balance hypothesis of obesity that emphasizes a calorie-is-a-calorie perspective [29–31]. Such a conclusion may be premature given the robust increase in EE associated with consumption of the KD as measured using doubly-labeled water in Hall et al. [8]. Indeed, on the basis of the increase in EE_DLW_ and, especially, in EE_nonchamber_, the results are entirely consistent with the carbohydrate-insulin hypothesis, which predicts an increase in EE with restriction of carbohydrate intake and the resulting decrease in insulin secretion. Overnight insulin withdrawal in patients with type 1 diabetes increases basal (resting) energy expenditure, a response that has been attributed to hyperglucagonemia and is associated with increased protein catabolism [see 32 for a review]. Although the chronic reduction in insulin secretion during the KD period in the Hall et al. study [8] was not as great, it was also associated with increases in basal metabolic rate (as estimated from sleeping energy expenditures), plasma glucagon concentrations, and protein catabolism (as indicated from urinary nitrogen, urea and ammonia excretion). These factors, along with a possible increase in gluconeogenesis, deserve further exploration in future studies of the effect of ketogenic diets on energy expenditure.

## Acknowledgments

We thank Marc Hellerstein, Mitchell Lazar and Michael Tordoff for their thoughtful and helpful comments on earlier drafts of this paper, John Thyfault for helpful discussions and suggestions, and William Wong for helpful assistance with the analysis and interpretation of doubly labeled water and respirometry data.

## Authors’ contributions

MF designed the analysis plan; SA performed the statistical analyses; MF and SA wrote the paper; MF had primary responsibility for final content.

## Supporting information

**S1 File. Data sources and handling (S1_file.pdf).**

**S2 File. SAS code for secondary analysis (SI_file.txt).**

**S3 File. Hall et al. 2019: Evaluation of Outlier Selection S3_file.pdf).**

## Supporting Information File S1

### Data Sources and Handling

#### Abbreviations

EE_chamber_, total daily energy expenditure measured in metabolic chambers; EE_DLW_, average energy expenditure measured by doubly labeled water; EE_nonchamber_, average energy expenditure on days subjects were living on the ward outside metabolic chambers measured by doubly labeled water; KD, low-carbohydrate/high-fat ketogenic diet; SEE, sleeping energy expenditure.

**Data Sources and Handling** (see Flowchart below for additional information)

#### Replication of calorimetry data

Data from Hall et al.’s Table 2 [1] for re-calculation and analysis of were extracted from the “Intake,” “chamber,” “BC” (body composition) and “DLW” (doubly labeled water) tabs of the Hall et al. dataset published on the Open Science Framework website [2]. In keeping with the original analyses, *P* values were not corrected for multiplicity in this replication or in the analyses below, which may limit inferences when multiple comparisons are made. Similarly, due to the small sample size, no multivariable model was applied to adjust for potential confounders.

#### Non-chamber EE_DLW_

Data to calculate Equation (1) and all other analyses of EE_DLW_, EE_nonc_ham_ber_, and EE_chamber_ were taken from the “DLW” tab in the Hall et al. dataset. EE_DLW_ values in Equation 1 correspond to the “TEE DLWChamber unadjusted” values in the “DLW” tab, which were derived using respiratory quotient measured in the chambers during the EE_DLW_ measurement periods. EE_chamber_ values in the “DLW” tab correspond to the “EE binned” values in the “Chamber” tab of the dataset averaged over the four chamber days during the 14-day EE_DLW_ measurement periods.

EE_nonchamber_ for each participant was also calculated using the difference between CO2 production rates measured in the chambers and by doubly labeled water (*rCO_2nonchamber_*) according to the following equation:

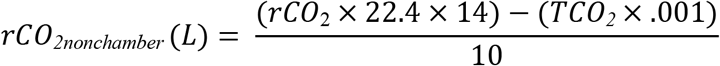

Where *rCO_2_* is the daily production rate of CO_2_ (mol/d), corresponding to “rCO_2_Redo values in the “DLW” tab of the Hall et al. dataset, which is multiplied by the molar volume of gas (22.4) and the number of days in the DLW measurement period. *TCO_2_* is the total CO_2_ production (in ml) measured by respirometry during the four days participants were housed in chambers, which were multiplied by 0.001 to convert to liters. These CO_2_ values were derived from “TVCO2” data in the “chamber” tab of the data set. The product of these calculations was divided by 10 to determine the daily volume of rCO_2_ (in L) produced during the 10 non-chamber days.

To calculate EE_nonchamber_ (kcal/d) during the BD period we used Equation 4 from Hall et al. [1] substituting *rCO_2nonchamber_* for *rCO_2_* as below:

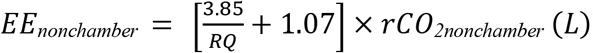 We used Hall et al.’s [1] Equation 5, which corrects for ketone body excretion (*K*_excr_ taken from “ketone_excr” values in the “DLW” tab of the dataset), to calculate EE_nonchamber_ during the KD period:

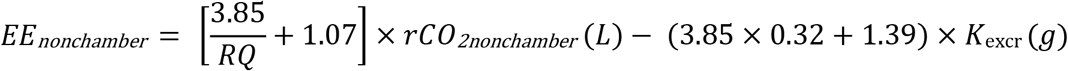

*RQ* is respiratory quotient and for both equations corresponds to the “RQ_Chamber” values in the “DLW” tab of the dataset.

#### EE_DLW_ outlier

For Subject 04-012, the first of the two body weight measurements associated with body composition assessments during the KD period was performed the day before the “Dose Date” for doubly labeled water (as indicated in the dataset “DLW” tab). The second was performed 12 days later, two days before the end of the EE_DLW_ measurement period. Body weights taken during body composition assessment (“BodyMass_kg”) were extracted from the “BC” (body composition) tab of the dataset and, for the KD EE_DLW_ measurement periods, from “DailyBW” tab of the dataset. Energy intake data were taken from the “EI” tab in the dataset.

#### Accelerometer data

Accelerometer counts were extracted from the “Accelerometer” tab in the dataset using the “Dose Date” for doubly labeled water in the dataset “DLW” tab as the first day of the EE_DLW_ measurement periods. We included data only from those days during which accelerometer wear time exceeded 720 minutes (12 hours) as specified in the Hall et al. code for analysis of the fractional difference in counts during chamber and non-chamber days.

#### Sleeping energy expenditure

SEE data were extracted as “SMR Chamber unadjusted” values from the “DLW” tab of the Hall et al. dataset. These values correspond to the “SMR binned” values in the “Chamber” tab of the dataset averaged over the chamber days during the EE_DLW_ measurement periods described above. As in Hall et al. [1], SEE (as kcal/d) was extrapolated from sleeping metabolic rate (as kcal/min).

---

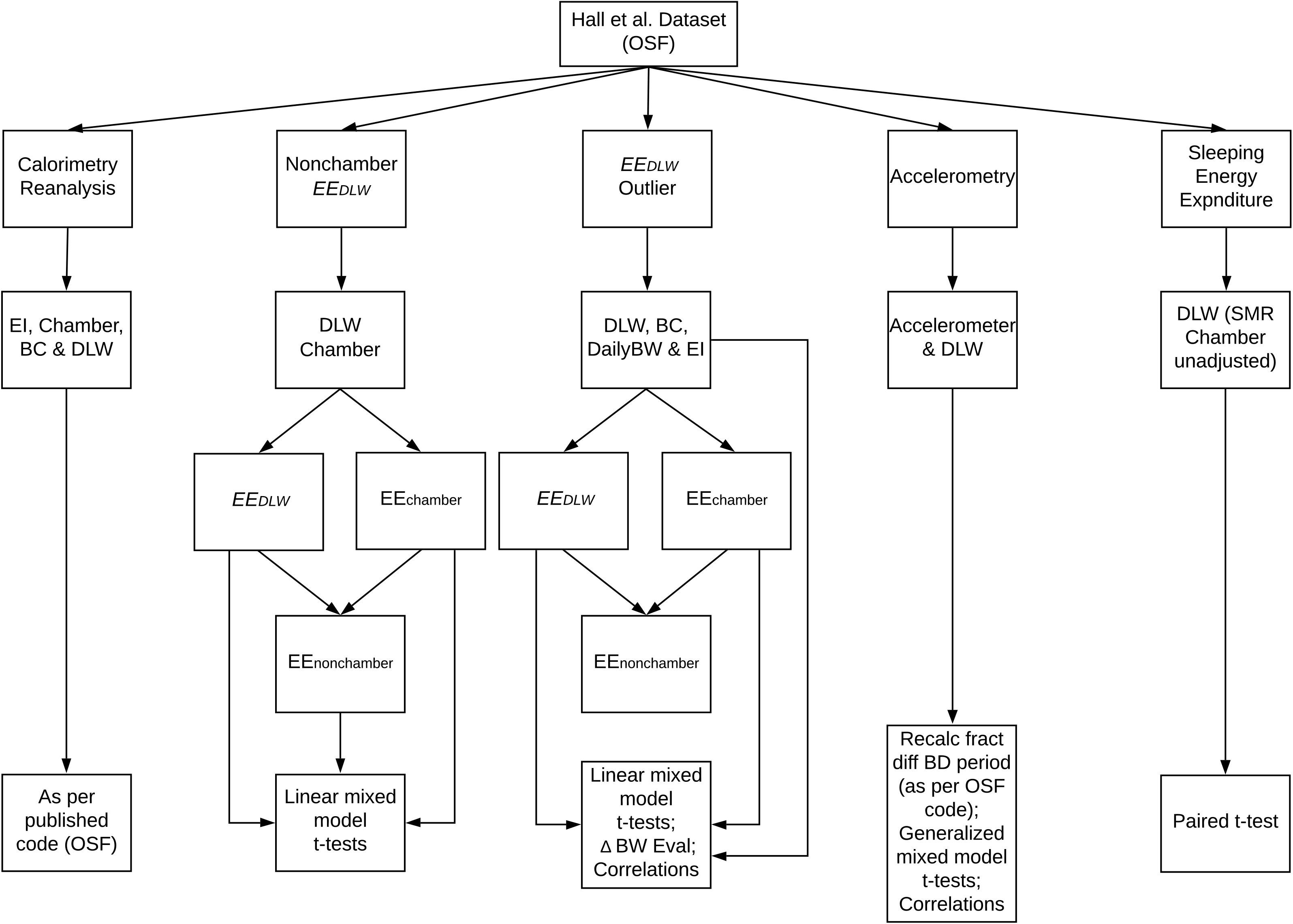

## Supporting Information File S2

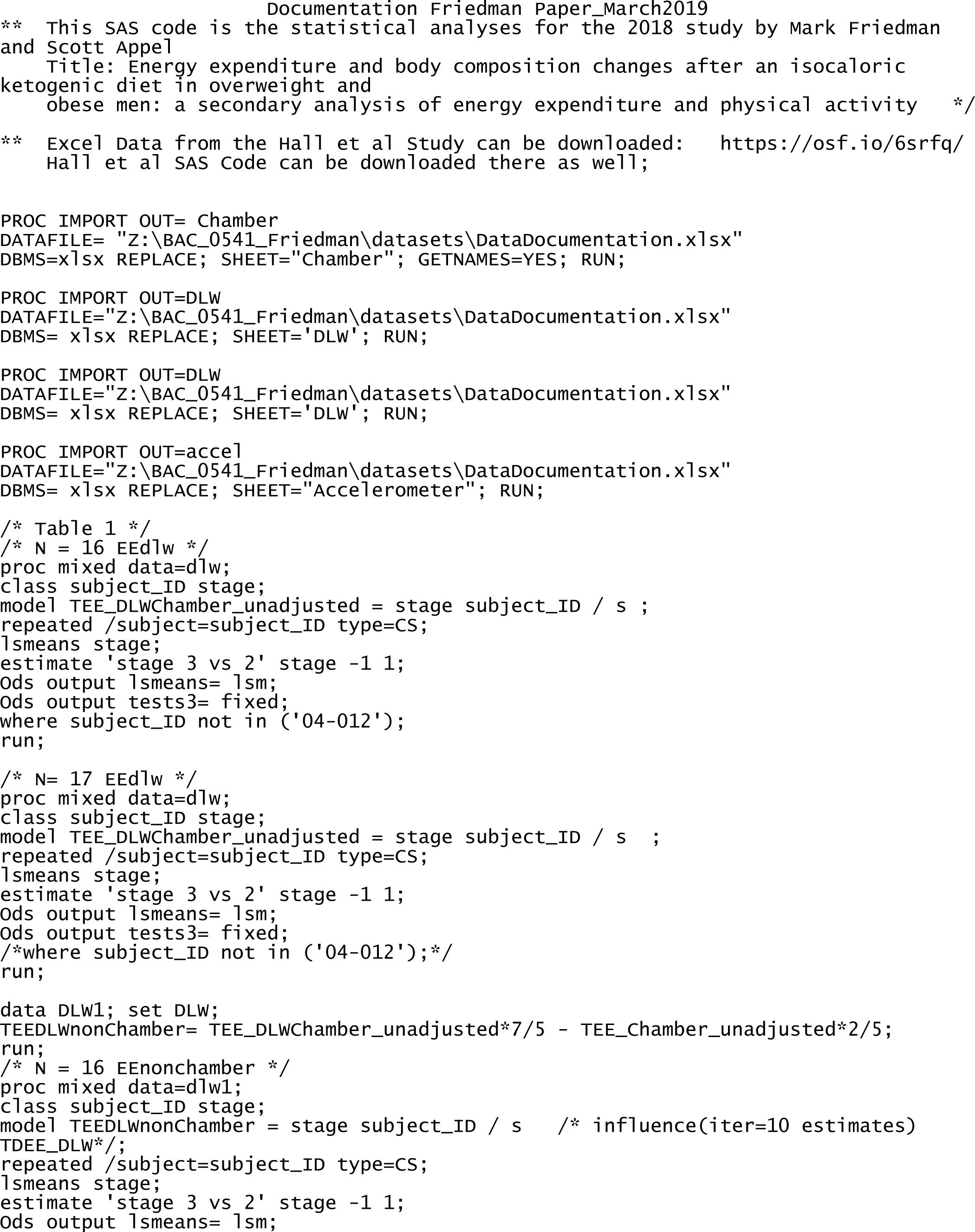
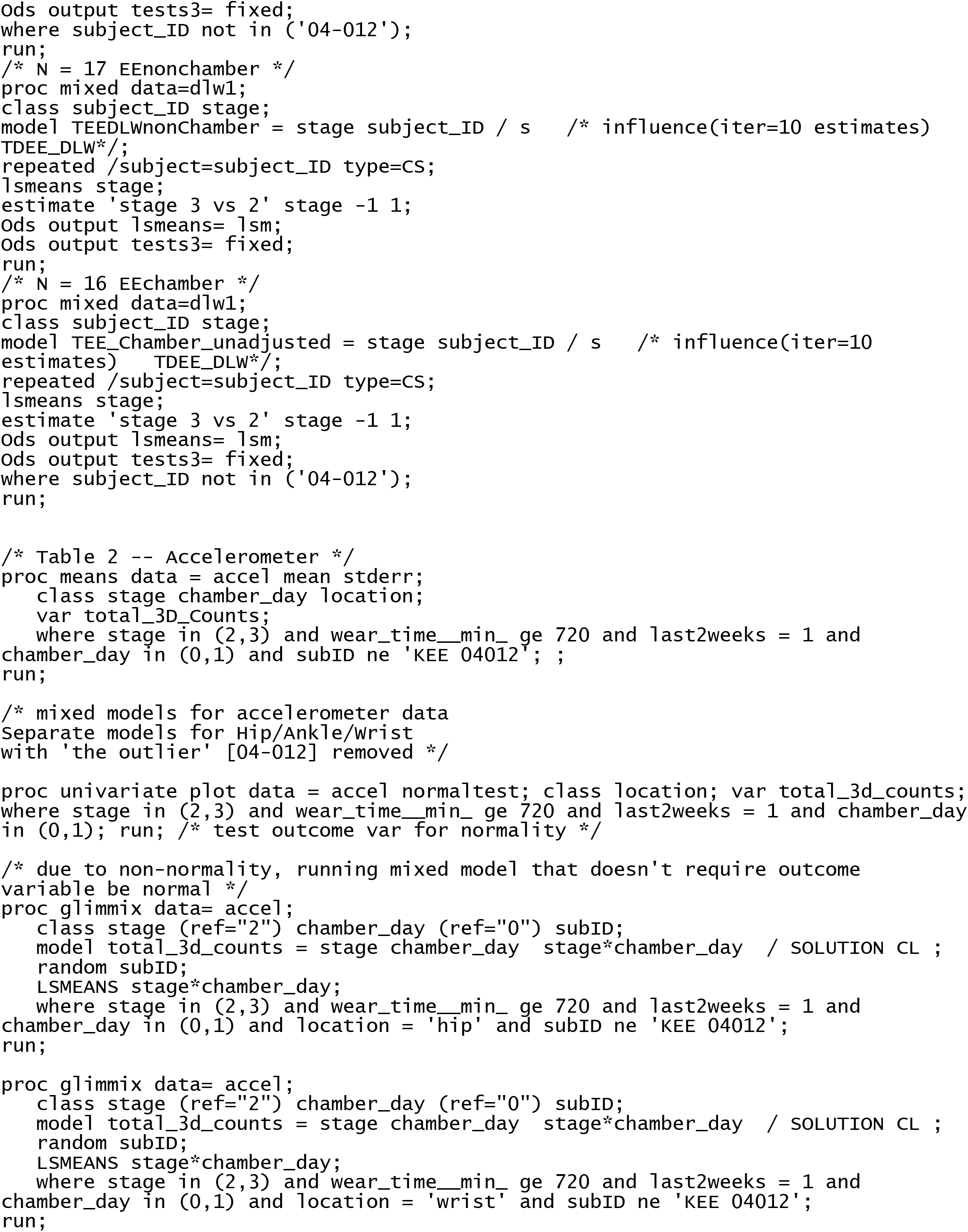
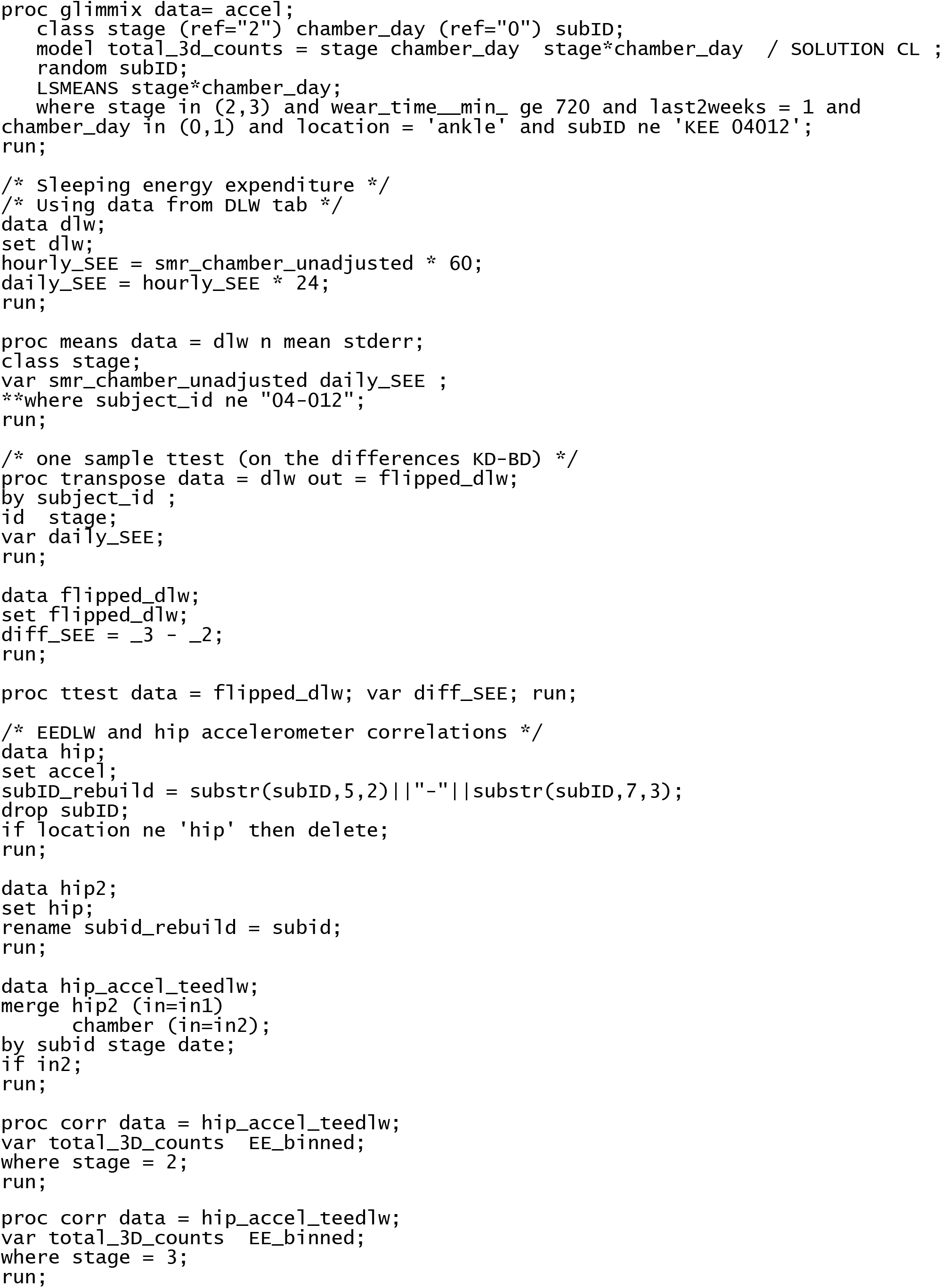

## Supporting Information File S3

### Hall et al. 2019: Evaluation of Outlier Selection

#### Restricted Selection

The protocol for the original Hall et al. study [1], which is the subject of their recent analysis [2], did not specify criteria or methods for identification and handling of outliers. In both studies, subjects were considered outliers, and their energy expenditure data excluded from analysis, based on *post hoc* observations indicating that their energy expenditure measured using doubly labeled water (EE_DLW_) was not commensurate with other parameters of energy balance. In their recent reanalysis of the original study, two outliers so identified were reported to have gained weight during the ketogenic diet (KD) period despite EE_DLW_ in excess of energy intake (EI). One of the two participants (“Subject A”) also gained weight during the BD period although EE_DLW_ exceeded energy intake and showed a “sleight” gain in fat mass during both periods. The other participant (“Subject B”) was also identified statistically as an outlier with respect to the magnitude of his increase in EE_DLW_ after the switch from the BD to KD diet. We examined the database for other participants who showed changes in body weight that were discrepant with respect to the difference between their EE_DLW_ and EI because this was a criterion for outlier status that was met by both Subject A and B.

Subjects A and B (Subject ID #’s 04-006 and 04-012, respectively; Group 1 in Table 1S) showed the greatest increase in EE_DLW_ over EI associated with weight gain of all 17 study participants.

**Table 1S.** Putative outliers based on discrepancies between body weight change and the difference in energy expenditure and energy intake.

| Group | ID# | $EE_{DLW}$<br>(kcal/d) | EI<br>(kcal/d) | $EE-EI$<br>(kcal/d) | $\Delta BW$<br>(kg) |
| --- | --- | --- | --- | --- | --- |
| 1 | 04-006<sup>a</sup> | 4448 | 2697 | 1751 | 0.6 |
|  | 04-012<sup>b</sup> | 3612 | 2794 | 818 | 0.2 |
| 2 | 02-004 | 2858 | 2393 | 465 | 1.3 |
|  | 03-009 | 2395 | 2645 | -250 | -0.2 |
|  | 04-007 | 3859 | 4150 | -291 | -1.9 |
| 3 | 03-002 | 2390 | 2445 | -55 | -0.4 |
|  | 03-008 | 2691 | 2745 | -54 | -0.2 |

<sup>a</sup> and <sup>b</sup> = Subjects A and B, respectively. All measures are during the ketogenic diet (KD) period. $EE_{DLW}$ and EE, energy expenditure by doubly labeled water; EI, energy intake; $\Delta BW$ , change in body weight during body composition assessments.

However, five other participants had changes in body weights during the KD period that were incommensurate with the difference between their EE_DLW_ and EI. Three participants (Group 2; Table 1S) showed more moderate, but substantial, differences between EE_DLW_ and EI. In two of these, body weight decreased despite an EE_DLW_ that was less than their EI, whereas the third gained weight although EE_DLW_ exceeded EI. Two additional participants (Group 3; Table 1S), exhibited small, negative differences between EE_DLW_ and EI that were associated with a decrease in body weight.

Subjects A and B showed the two greatest increases in EE_DLW_ and nonchamber energy expenditure (EE_nonchamber_) after the switch from the BD to KD of all 17 participants. Therefore, exclusion of their expenditure data would be expected to reduce any increase in average expenditures after the switch whether adjusted for energy balance or not. Figure 1S shows the effect of excluding these two participants (Group 1) on average EE_nonchamber_, a primary outcome of our analysis, along with the effect of excluding other outliers (Groups 2 and 3) listed in Table 1S either separately or in combination with those in Group 1.

**Figure 1S.**
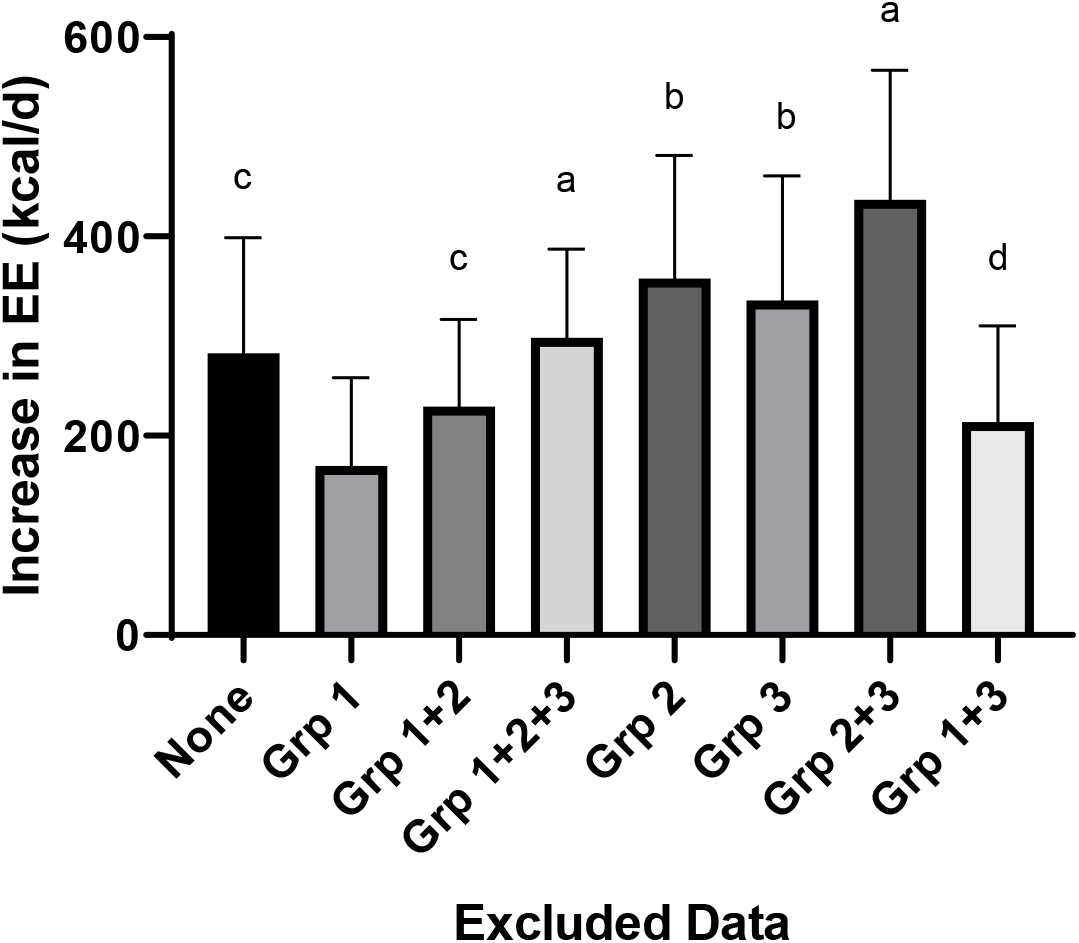
Effect of excluding putative outlier data on the increase in non-chamber energy expenditure by doubly labeled water (EE_noncharaber_) after switching from a basal to a ketogenic diet. Values are mean ± SE. a, b, c, and d = *P* < 0.01, 0.02, 0.03, and 0.05, respectively, by paired t-test. For Group 1, *P* = 0.076.

Although EE_nonchamber_ increased significantly after the switch to the KD when all 17 participants are included in the analysis, the increase was not statistically significant when data from Subjects A and B (Group 1) were excluded from the analysis. In contrast, the increase of EE_nonchamber_ was significant despite removal of other groups of putative outliers either alone or in combination with other groups. The effect of excluding Group 2 alone or in combination with Group 1 is especially notable because the energy imbalances reflected by the differences between EE_DLW_ and EI, while less than that in Group 1, were substantial. As reported in this paper (Table 1), the increase in EE_nonchamber_ after the diet switch was statistically significant after exclusion of Subject B’s (ID# 04-012) data from the analysis. When only Subject A’s data was removed from the analysis, the increase in EE_nonchamber_ after the diet switch (257 ± 116 kcal/d) was also statistically significant (*P* =0.49 by paired t-test).

#### Asynchronous Measurement Intervals

The reported gain of body weights during the KD period of Subjects A and B was based on body weight measurements taken during two body composition assessments in the latter part of the diet period. Hall et al. [2] claimed that the interval for body composition assessments was coincident with the EE_DLW_ measurement period. However, inspection of dates in the original study’s dataset for DLW dosing and body composition assessments shows this was not the case; body composition and EE_DLW_ measurements periods were coincident in only 6 and 4 out of 17 participants in the BD and KD periods, respectively.

Because the dataset includes dates and daily body weight measurements for all subjects throughout the study, it is possible to determine the change in body weight over the EE_DLW_ measurement period independently from the body weight measures taken during body composition assessments.

As discussed in this paper, Hall et al. [1] reported that Subject B gained 0.2 kg during the KD EE_DLW_ measurement period based on body composition assessments (Table 2S), but daily body weight measurements show a body weight loss of 0.5 kg during the actual EE_DLW_ measurement period (Table 2S). Similarly, according to the database, Subject A gained 0.6 kg of weight as per body composition assessments, but according to daily body weight measurements, lost 0.5 kg during the EEDLW measurement period.

**Table 2S.** Changes in body weight in putative outliers as a function of measurement interval during the ketogenic diet period.

|  |  | **BC Period** | **BWBC (kg)** | **BWCD (kg)** | **DLW period** | **BWDD (kg)** |
| --- | --- | --- | --- | --- | --- | --- |
| **Subject A** | Start | 26-Mar-14 | 63.9 | 64.5 | 28-Mar-14 | 64.6 |
|  | End | 9-Apr-14 | 64.5 | 65.0 | 10-Apr-14 | 64.1 |
|  | **diff** | -- | **0.6** | **0.5** | -- | **-0.5** |
| **Subject B** | Start | 18-Jun-14 | 88.6 | 89.1 | 19-Jun-14 | 89.0 |
|  | End | 30-Jun-14 | 88.8 | 89.1 | 2-Jul-14 | 88.5 |
|  | **diff** | -- | **0.2** | **0.0** | -- | **-0.5** |

**BC**, body composition assessment dates; **DLW**, doubly labeled water measurement dates; **BWBC**, body weight for body composition measurement dates using body composition assessment dates; **BWCD**, body weight for body composition measurement dates using daily body weight measurements; **BWDD**, body weight for doubly labeled water measurement dates using daily body weight measurements; **diff**, change in body weight from start to end.

No other participants showed such a reversal of body weight change from gain to loss (or vice versa) during the KD period when body weight measurements were synchronized with the EE_DLW_ measurement period, although one participant did so during the BD period. The changes in body weight of Subjects A and B over the EE_DLW_ measurement period measured using daily body weight data were well within one standard deviation of that for the group as a whole (−0.9 ± 0.6.kg, mean ± SD). The differences between body weight measurements from body composition assessments and recorded daily body weights do not appear to be due to differences in the precision of measurement under the two conditions because changes in body weight over the two body composition assessments as determined using daily body weight measurements were consistent with those measured during composition evaluations (Table S2).

